# Neonicotinoid and pyrethroid combination: A tool to manage insecticide resistance in malaria vectors? Insights from experimental evolution

**DOI:** 10.1101/2021.06.09.447494

**Authors:** Marius Gonse Zoh, Jean-Marc Bonneville, Jordan Tutagana, Frederic Laporte, Behi K. Fodjo, Chouaibou S. Mouhamadou, Christabel Sadia, Justin McBeath, Frederic Schmitt, Sebastian Horstmann, Stephane Reynaud, Jean-Philippe David

## Abstract

**Background:** The introduction of neonicotinoids for managing insecticide resistance in mosquitoes is of high interest as they interact with a biochemical target not previously used in public health. In this concern, Bayer developed a combination of the neonicotinoid clothianidin and the pyrethroid deltamethrin (brand name Fludora® Fusion) as a new vector control tool. Although this combination proved to be efficient against pyrethroid-resistant mosquitoes, its ability to prevent the selection of pyrethroid and neonicotinoid resistance alleles was not investigated. In this context, the objective of this work was to study the dynamics and the molecular mechanisms of resistance of *An. gambiae* to the separated or combined components of this combination. A field-derived *An. gambiae* line carrying resistance alleles to multiple insecticides at low frequencies was used as a starting for 33 successive generations of controlled selection. Resistance levels to each insecticide and target site mutation frequencies were monitored throughout the selection process. Cross resistance to other public health insecticides were also investigated. RNA-seq was used to compare gene transcription variations and polymorphisms across all lines.

**Results:** This study confirmed the potential of this insecticide combination to impair the selection of resistance as compared to its two separated components. Deltamethrin selection led to the rapid enrichment of the kdr L1014F target-site mutation while clothianidin selection led to the over-transcription of multiple cytochrome P450s including some showing high homology with the ones conferring neonicotinoid resistance in other insects. A strong selection signature associated with clothianidin selection was observed on a cytochrome P450 gene cluster previously associated with resistance. Within this cluster, the gene *CYP6M1* showed the highest selection signature together with a transcription profile supporting a role in clothianidin resistance. Modelling the impact of point mutations selected by clothianidin on CYP6M1 protein structure suggested that the selection of variants affecting its active site can enhance clothianidin metabolism.

**Conclusions:** In the context of the recent deployment of neonicotinoids for mosquito control and their frequent usage in agriculture, the present study highlights the benefit of combining them with other insecticides for preventing the selection of resistance and sustaining vector control activities.

## Introduction

Malaria, caused by the protozoan *Plasmodium* parasite and transmitted by Anopheles mosquitoes is a major public health problem in Africa with 213 million cases and 405 000 deaths in 2018 [1]. Without effective vaccines and the development of antimalarial drug resistance by parasites, the fight against malaria mainly relies on vector control operations using insecticides [2] through insecticide-treated nets (ITNs) and indoor residual spraying (IRS) [3,4]. Among the insecticides used in public health, pyrethroids remain the most used on ITNs because of their strong effects against the target insects and their low toxicity to the environment and mammals. In addition to pyrethroids, other insecticide families such as carbamates, organophosphates and in some specific cases DDT are also used in public health to improve the effectiveness of the fight against malaria vectors [5,6].

However, malaria vector control is now threatened by the spread of pyrethroid resistance throughout Africa [7]. Resistance to pyrethroids can be the consequence of various mechanisms in mosquitoes, such as non-synonymous mutations affecting the voltage-gated sodium channel targeted by these insecticides (i.e. Knock Down Resistance ‘kdr’ mutations), a lower insecticide penetration through insect cuticle, its sequestration, or its biodegradation (metabolic resistance) [8,9]. *Kdr* mutations can confer resistance to pyrethroids and DDT. These mutations prevent the insecticide from binding to the voltage-gated sodium channel (VGSC) of the insect’s nervous system. *Kdr* mutations affecting malaria vectors are widely distributed in Africa with two distinct mutations at position 1014 of the VGSC mainly associated with insecticide resistance [10,11]: the L1014F *kdr west* mutation and the L1014S *kdr east* mutation [12,13]. Metabolic resistance is also widespread in African malaria vectors and has been associated with resistance to multiple insecticide families. Increased insecticide metabolism can be caused by an increased activity of detoxification enzymes or structural modification affecting insecticide turnover. These detoxification enzymes include cytochrome P450 monooxygenases (P450s or *CYP* for genes), carboxy/cholinesterases (CCEs), glutathione S-transferases (GSTs) and UDP-glycosyl-transferases (UDPGTs) although other families can be involved [9,14,15]. P450s from the *CYP6* gene family and epsilon GSTs have been mainly associated with metabolic resistance to pyrethroids in *An. gambiae* [14]. Overall, though target-site modifications and metabolic resistance appear to play a central role in pyrethroid resistance, additional resistance mechanisms such as cuticle modification, altered transport and sequestration have also been reported [16–18].

Most African *An. gambiae* populations are resistant to pyrethroid insecticides and also show varying levels of resistance to other insecticides used for vector control (carbamates, organophosphates and organochlorines). Insecticide resistance management therefore will benefit from the introduction of new chemistries. The fastest way to achieve that is by repurposing chemicals already used in agriculture, especially those with different modes of action which are less likely to be affected by cross-resistance with current public health insecticides [19]. Among the insecticides which are being proposed for managing pyrethroid resistance, neonicotinoids have aroused a high interest as i) they have not previously been used for public health, ii) they show a good toxicity against mosquitoes and a low toxicity to mammals [20] and iii) they target nicotinic acetylcholine receptors (nAChR) which constitute a novel target for malaria vectors [21,22]. In this situation, a new insecticide formulation combining the neonicotinoid clothianidin and the pyrethroid deltamethrin (8:1 w/w) under brand name *Fludora® Fusion* was developed by Bayer to be an additional tool for insecticide resistance management in African malaria vectors. By combining two unrelated modes of action, this indoor residual spraying (IRS) combination is intended to slow down the selection of resistance alleles to the newly introduced neonicotinoid clothianidin. Field trials suggested that *Fludora® Fusion* shows a good residual activity against malaria vectors from different African countries [23,24] and a good efficiency against pyrethroid-resistant *An. gambiae* [25]. Despite its efficacy, the hypothesis that a combination of two modes of action offers a different selection profile, or indeed a benefit, as compared to the individual insecticide components has not been tested in malaria vectors. As neonicotinoids are often used against crop pests in Africa, resistance alleles to this insecticide family may already be circulating at low frequency in natural mosquito populations neighboring agriculture areas. In such context, the large-scale implementation of *Fludora® Fusion* for vector control might lead to the rapid selection of these resistance alleles leading to a decreased efficacy of this novel formulation, particularly if pyrethroid resistance alleles are also present. Such hypothesis is supported by a recent study suggesting the presence of neonicotinoid resistance alleles in *Anopheles coluzzi* populations neighboring agriculture areas [26,27].

In this context, the primary objective of this work was to compare the dynamics of resistance between the combination *Fludora® Fusion* and its two individual insecticide components in *An. gambiae* and assess the ability of the combination to hinder resistance selection. The secondary objective of this work was to investigate the associated resistance mechanisms using molecular approaches. An *An. gambiae* line carrying resistance alleles to multiple insecticide families at low frequency was created by crossing a multi-resistant strain originating from an intense agricultural area of Côte d’Ivoire and a susceptible strain. The resulting line was then used for selecting, across multiple generations, three lines with *Fludora® Fusion*, deltamethrin alone or clothianidin alone. The evolution of resistance levels to each insecticide and target site mutation frequencies was monitored in each line throughout the selection process. Cross resistance profiles of each line to insecticides used for selection and to other public health insecticides were also investigated. A comparative RNA-sequencing approach was used to characterize the impact of selection on each line at the transcriptome scale. Both differential gene transcription levels and selection signatures based on polymorphism data were investigated. The role of particular P450s in clothianidin resistance were then further explored using RT-qPCR and *in silico* protein modeling. The outcomes of the present study are discussed in the context of the deployment of neonicotinoids for malaria control and the added value of insecticide combinations for insecticide resistance management.

## Results

### Resistance dynamics during the selection process

WHO susceptibility assays confirmed the low frequency of resistance alleles to most insecticides in the Tiassalé-S line with mortality levels higher than 95% for the insecticides deltamethrin, bendiocarb and fenitrothion (Additional file 1). However, despite the introgression of susceptible alleles by controlled crosses, the frequency of resistance alleles to DDT was still high with only 46% mortality observed to 4% DDT. Monitoring the resistance of each selected line to its respective insecticide along the selection process showed a rapid rise of deltamethrin resistance in the Delta-R line as compared to the non-selected line Tiassalé-S (Figure 1). In the Delta-R line, mortality following deltamethrin exposure dropped from 73% at G0 to 30% at G7 and further decreased in the following generations to stabilize around 15% (P<0.05 from G7 to G33). In the Clothia-R line, a significant increased resistance to clothianidin was observed from generation G13 as compared to the non-selected line (P<0.05 from G13 to G33) with resistance stabilizing at a moderate level. In the Fludo-R line, a significant increased resistance Fludora® Fusion mixture was observed at G13 (P<0.05) though this trend was not confirmed in the subsequent generations.

**Figure 1.**
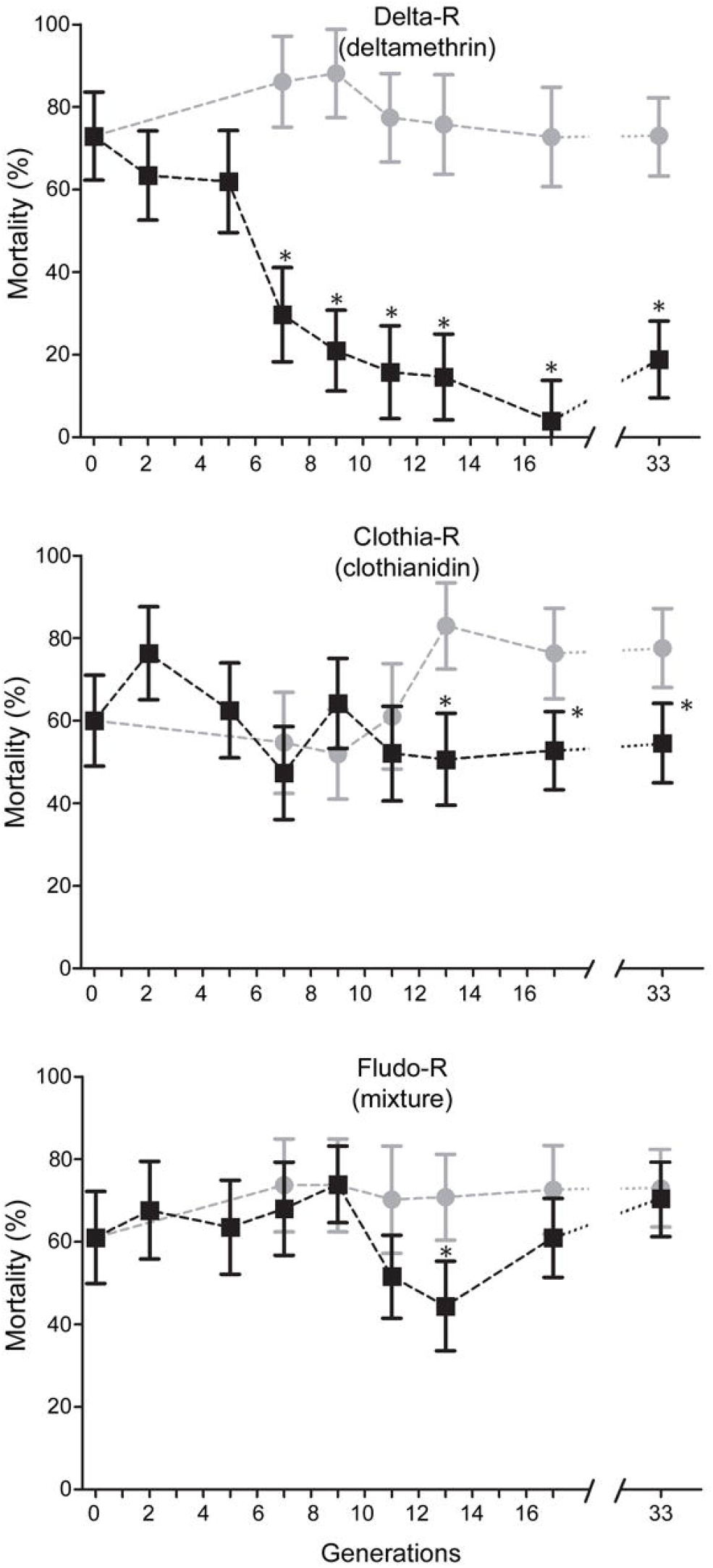
Resistance dynamics of each line along the selection process. Black squares: resistance of each selected line to the insecticide used for selection. Grey dots: resistance of the non-selected Tiassalé-S line. Resistance levels were compared using bottle assays and are expressed as mean % mortality ± 95% Wald confidence interval. Comparisons between each line and the Tiassalé-S line at each time point were performed using a Fisher test on mortality proportions (* p<0.05).

Assessing the cross-resistance of each line to insecticides used for selection at G17 confirmed the high resistance of the Delta-R line to deltamethrin (P<0.05) but also indicated a slight increased resistance of the Fludo-R line to deltamethrin (P<0.05) (Figure 2A). A slight increased resistance to clothianidin was observed for the Clothia-R and Fludo-R lines (P<0.05). Delta-R and Clothia-R lines showed a slight increased resistance to Fludora® Fusion mixture (P<0.05) while no such increased resistance was observed for Fludo-R line (P>0.05). Assessing the cross-resistance of each selected line to other insecticides used in public health confirmed the moderate resistance of the Tiassalé-S line to DDT and further indicated that DDT resistance alleles were further enriched in all selected lines with mortality levels dropping to less than 10% (P<0.05) (Figure 2B). No significant increased resistance to fenitrothion or bendiocarb was detected for any selected line.

**Figure 2.**
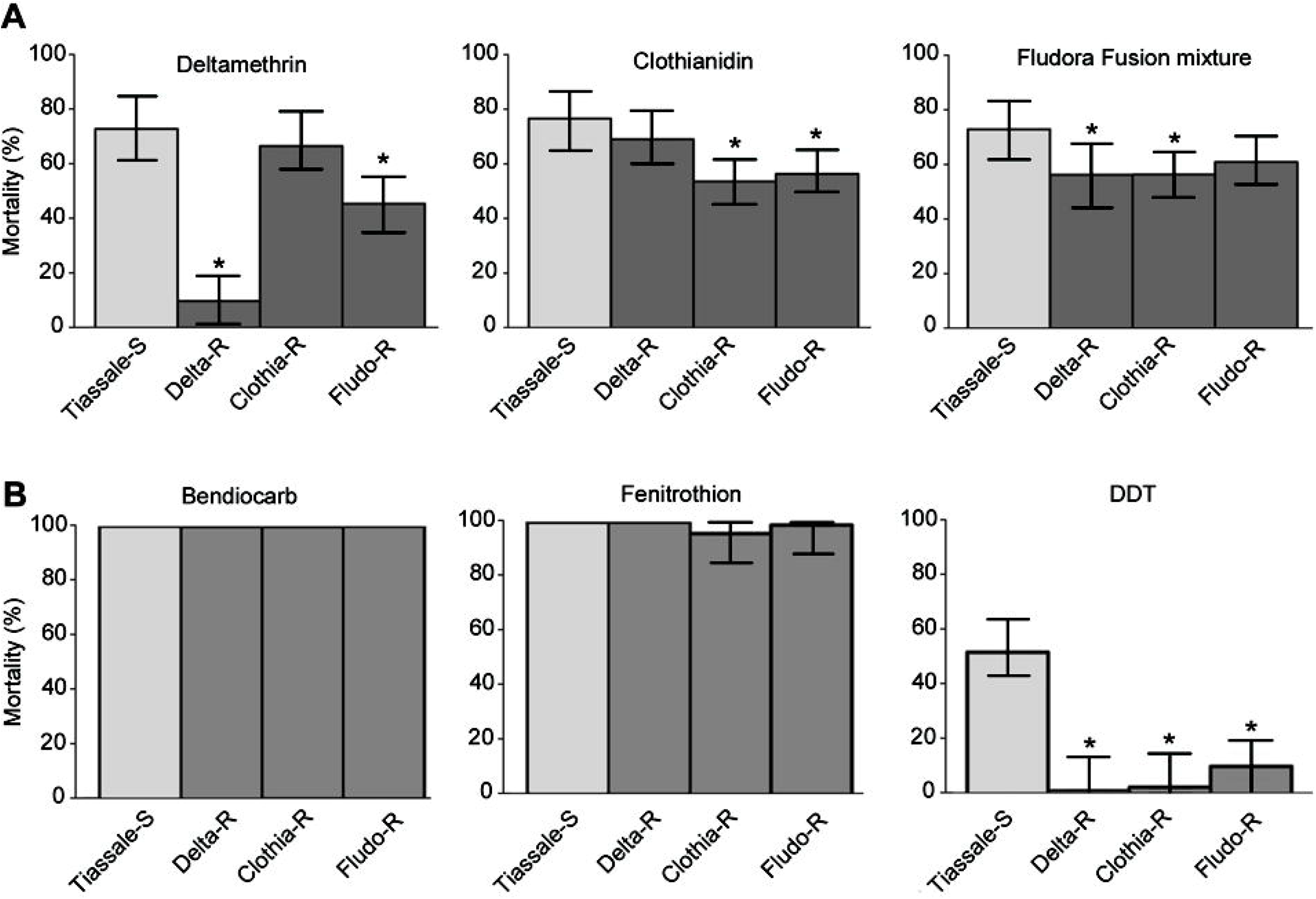
Cross-resistance of each line to insecticides. Cross-resistance profiles of each line to insecticides used for selection (A) and to other insecticide families used for vector control (B). Resistance levels to the insecticides used for selection were compared using bottle assays while resistance levels to other insecticides were compared using WHO test tubes equipped with papers impregnated with 0.5% bendiocarb, 1% fenitrothion and 4% DDT. Mortality rates are expressed as mean % mortality ± 95% Wald confidence interval and were compared to the unselected line using a Fisher test on mortality proportions (* p<0.05).

### Target sites mutations

The genotyping of the kdr west mutation (L1014F) revealed its association with deltamethrin resistance in the Delta-R line with an increased frequency from 47% at G0 to nearly 100% at G7 with most individuals being homozygotes for the resistant allele 1014F (Figure 3). Although fluctuations were observed during the selection process in the Clothia-R and Fludo-R lines, the frequency of the resistant allele did not increase in response to selection with clothianidin or Fludora® Fusion mixture. A gradual decrease of resistant homozygotes was even observed in both lines. As expected because not targeted by any insecticide used for selection, the frequency of the G119S Ace1 mutation was not affected by insecticide selection in any line with an estimated frequency of 10%, 6%, 15% and 10% in the Tiassalé-S line (G0), Delta-R line (G17), Clothia-R line (G17) and Fludo-R line (G17) respectively (not shown).

**Figure 3.**
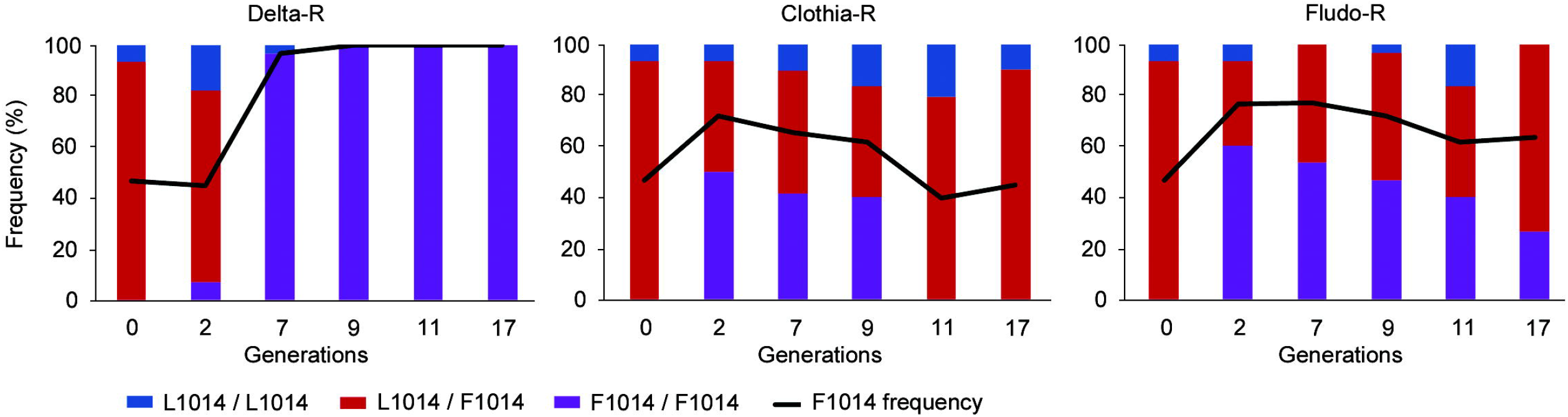
Dynamics of the L1014F kdr mutation along the selection process. For each line, the L1014F kdr mutation was genotyped in individual mosquitoes using the allele-specific qPCR TaqMan assay described in Bass et al. (2010). For each selected line, both genotype and allele frequencies are reported.

### Gene transcription levels

RNA sequencing produced >120M reads across four replicates for each line. Though ∼10% of these reads were filtered out based on sequence and mapping quality, this allowed detecting transcription signal in adult females for 10,839 genes. Among them, 2,628 genes were found differentially transcribed in at least one selected line as compared to the non-selected Tiassalé-S line (FC ≥1.5-fold and adjusted P value ≤0.005, Additional file 2). A higher number of genes were differentially transcribed in the Delta-R (1440 genes) and Clothia-R (1975 genes) lines as compared to the Fludo-R line (463 genes) (Additional file 3). Only 31 and 17 genes were found over- and under-transcribed in all selected lines respectively.

Gene Ontology (GO) term enrichment analysis performed on genes significantly over-transcribed in selected lines revealed a significant enrichment of terms associated with P450 activity in the Clothia-R line and in a lesser extent in the Fludo-R line (Additional file 4). These included the terms ‘monooxygenase activity’, ‘oxidoreductase activity, acting on paired donors, with incorporation or reduction of molecular oxygen’, ‘heme binding’ and ‘iron ion binding’. Terms associated with acetylcholine receptors were enriched in the Clothia-R and Delta-R lines including the terms ‘acetylcholine-activated cation-selective channel activity’ and ‘acetylcholine binding’. Enrichment analysis performed on genes significantly under-transcribed showed no enrichment of functions classically associated with resistance except a slight enrichment of terms associated with P450 and esterase activities in the Fludo-R line.

Focusing on gene families associated with known resistance mechanisms revealed different gene transcription profiles in each selected line as compared to the non-selected Tiassalé-S line (Figure 4). Five P450s belonging to *CYP325, CYP6* and *CYP4* families, six esterases, two UDPGTs (*AGAP006775* and *AGAP009562*) and four cuticular proteins were over-transcribed in the Delta-R line. Among them only *CYP4J10* was previously shown to be over-transcribed in association with pyrethroid resistance. The Clothia-R line showed an over-transcription of multiple candidate genes belonging to families commonly associated with insecticide resistance. This included 16 P450s, most belonging to the *CYP6, CYP9, CYP12* and *CYP4* families. Some of them (*CYP6M1, CYP12F1, CYP4C27, CYP6P1, CYP4G16, CYP6M3, CYP6Z3, CYP6Z2, CYP4H24, CYP6Z1, CYP6P3, CYP6P4*) were previously associated with insecticide resistance in *An. gambiae* or showed a high protein similarity with P450s conferring neonicotinoid resistance in other insect species (see methods). Five carboxylesterases were also over-transcribed in the Clothia-R line, with *COEAE2G* having been previously associated with insecticide resistance. Nine transferases were also over-transcribed in the Clothia-R line, most of them belonging to the UDPGT family. In addition to detoxification enzymes, clothianidin selection was also associated with the over-transcription of six ABC transporters and six cuticular proteins among which *CPR21* and *CPAP3-A1b* were both previously associated with resistance. In addition, clothianidin and deltamethrin selection were both associated with the over-transcription of multiple nicotinic acetylcholine receptor subunits. Fewer candidate genes were over-transcribed in the Fludo-R line with most of them being shared with the Clothia-R line. This included four *CYP4*s (*CYP6AH1*, CYP*4D17, CYP4C17 and CYP4D15*), one esterase (*AGAP011509*), two sulfotransferases (*AGAP029784, AGAP029783*), one GST (*GSTD11*) and two ABC transporter (*ABCCA3, ABCG18*). Only one UDPGT gene (AGAP006775) was over-transcribed in the three selected lines as compared to the non-selected Tiassalé-S line.

**Figure 4.**
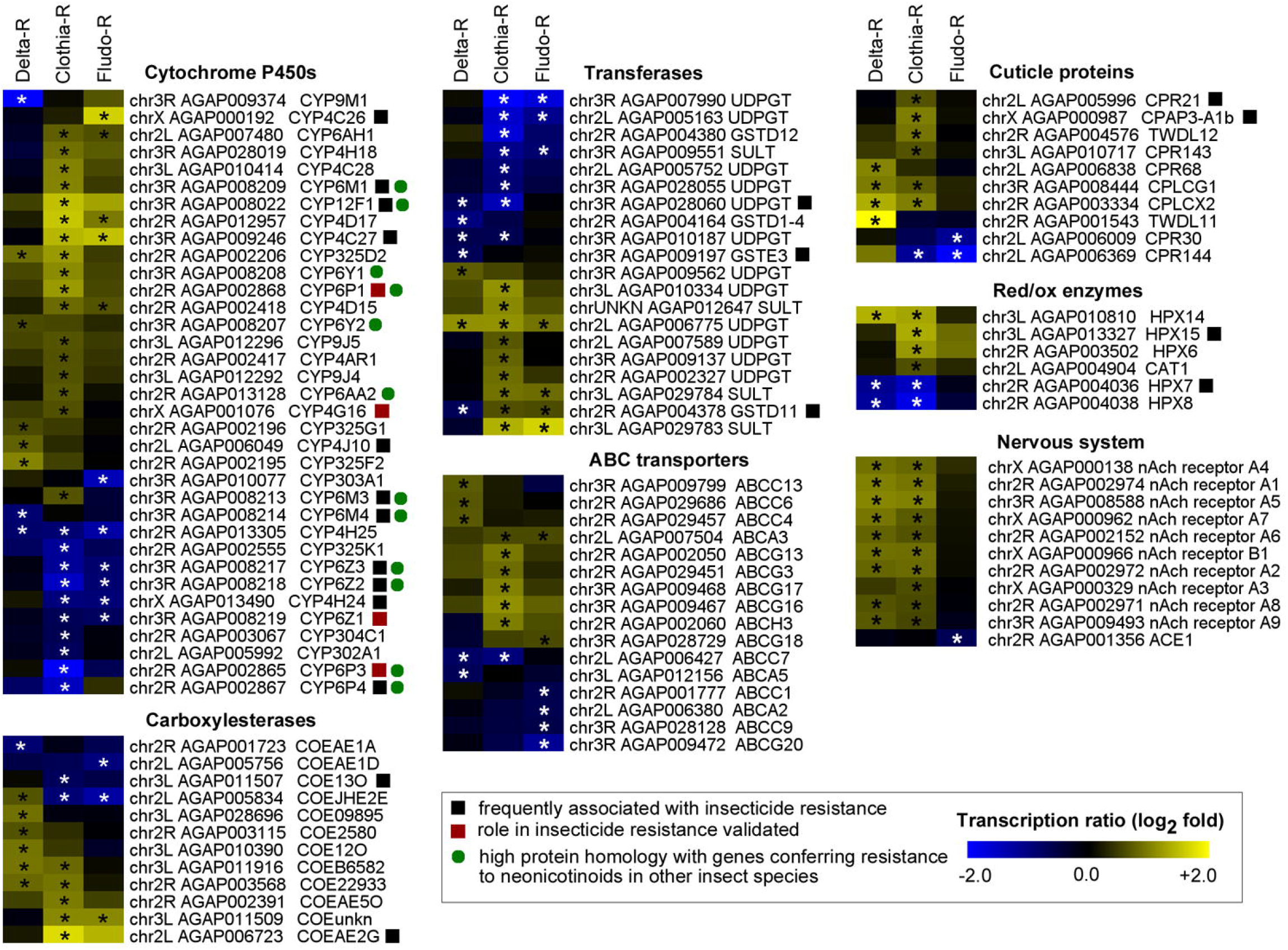
Transcription profiles of candidate genes associated with resistance. All candidate genes differentially transcribed in at least one selected line are shown. Colour scale shows the mean Log_2_ Fold Change between each selected line and the parental Tiassalé-S line. Stars indicate a significant differential transcription (FC>1.5-fold and corrected P value ≤ 0.005). Black squares indicate genes frequently associated insecticide resistance. Red squares indicate functionally validated resistance genes. Green dots indicate genes showing high protein homology with genes conferring resistance to neonicotinoids in other insect species.

### Polymorphism variations

RNA-seq allowed detecting 145008 bi-allelic substitutions or indels being polymorphic between the Tiassalé-S line and any selected line. These SNPs were mostly located within gene boundaries (>99%) and covered ∼44% of *An. gambiae* genes. Within genes affected by polymorphic variations, 50% of them carried at least 15 variations. These 145K variations led to >200K predicted genic effects according to AgamP4.12 annotation including ∼8.3% of them affecting protein sequence. Looking for selection signatures differentiating the Tiassalé-S line from each selected line based on allele frequency variations identified less differential SNPs in the Delta-R line (283, 0.19%) than in the Clothia-R line (2632, 1.79%) and the Fludo-R line (1397, 0.96%). A similar trend was also observed using the Bayesian FST-based approach also slightly more outliers were identified (from 0.95% to 1.95%). Summing up outliers/differential SNPs at the gene level and projecting them on the AgamP4 genome revealed different selection signature profiles between the three selected lines (Figure 5 and Additional file 5). Only a few loci showed strong selection signatures in the Delta-R line, none matching with gene previously associated with insecticide resistance. No selection signature was observed at the *Kdr* mutation locus though no polymorphic SNP was detected in this region due to the low transcription level of the *VGSC* gene. The closest polymorphic SNPs located ∼80 Kb upstream and ∼140 Kb downstream did not show any significant selection signature. Conversely, the Clothia-R line showed multiple loci showing strong selection signatures essentially located on chr 2R, 3R and 3L. The two loci located on chr 2R at ∼5.5 Mb and on chr 3L at ∼19.8 Mb were not associated with any candidate gene. The locus located on chr 3R at ∼6.9 Mb matched the *CYP6M* locus previously associated with insecticide resistance. Finally, the Fludo-R line showed some selection signatures but none of them were overlapping those observed in the two other lines. The two strongest selection signatures observed on chr 3R at ∼10 Mb and on chr 3R at ∼11 Mb did not match any candidate gene or known resistance locus.

**Figure 5.**
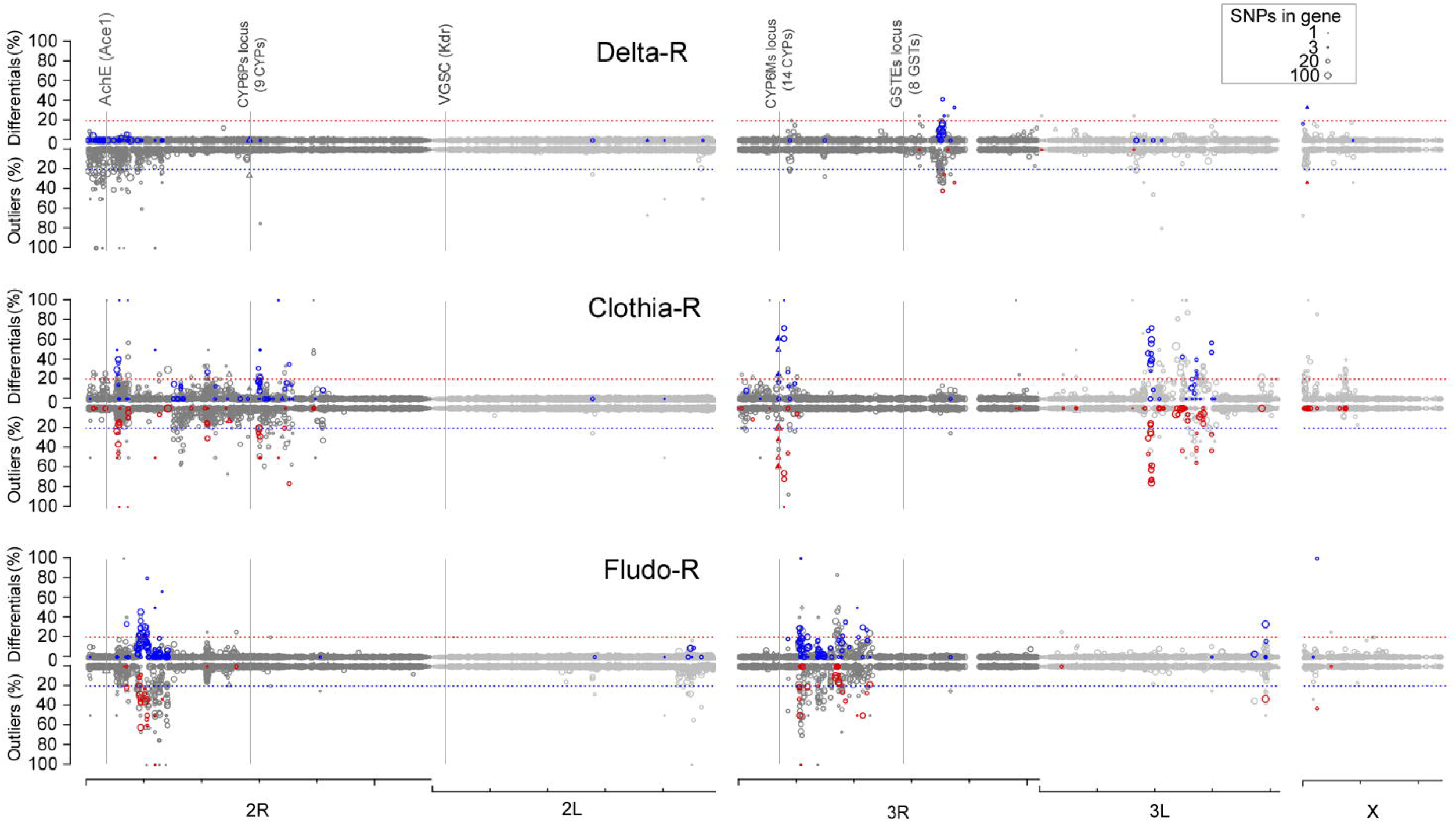
Selection signatures observed in each selected line. Selection signatures as compared to the non-selected Tiassalé-S line were computed using the 145008 bi-allelic polymorphic SNPs detected by RNA-seq. The upper Y axis shows the proportion of differential SNPs per gene as obtained by the frequency-based approach while the lower Y axis shows the proportion of outliers per gene as obtained by the Fst-based approach. For each approach, blue/red marks indicate genes showing a differential/outlier proportion higher than 20% with the alternative approach (red/blue dashed lines). Symbol size is proportional to the number of polymorphic SNPs per gene. Triangles indicate candidate genes potentially involved in insecticide resistance. Filled symbols indicate the presence of differential SNPs / outliers affecting protein sequence. Loci commonly associated with insecticide resistance in *An. gambiae* are indicated (VGSC, AchE, CYP6Ps cluster, CYP6Ms cluster and GSTEs cluster). The genomic scale shows chromosome arms with ticks every 10 Mb.

### Association of CYP6Ms with clothianidin resistance

The *CYP6M* locus showing a strong and specific selection signature in the Clothia-R line included 14 distinct *CYP6* genes (Figure 6A). In addition to *CYP6M2* known to metabolize pyrethroids for which no polymorphic SNP was detected, this locus contained 8 *CYP6* genes (*CYP6Y2, CYP6Y1, CYP6M1, CYP6M3, CYP6M4, CYP6Z1, CYP6Z2, CYP6Z3*) showing high protein homology with P450s previously associated with neonicotinoid resistance in other insects (see methods). Among them, *CYP6Y1, CYP6M1* and *CYP6M3* were all over-transcribed in the Clothia-R line. The strongest selection signature was detected for *CYP6M1* which contained 28 differential/outlier SNPs out of 44 with 11 of them being non-synonymous (Figure 6B) leading to the following amino acid changes: (G>A) V35I, (A>C) I66L, (G>A) G171D, (A>C) E215D, (T>G) F240L, (C>A) D347E, (C>G) D351E, (T>C) M375T, (G>T) A377S, (T>G) F396L, (T>G) S401A. Among them, E215D, F240L and M375T were located within CYP6M1 substrate recognition sites SRS2, SRS3 and SRS5 respectively (Figure 6C). In addition, E215D and F240L matched positions previously shown to interact with the binding of the neonicotinoid imidacloprid in *B. tabaci* CYP6CM1vQ, *D. melanogaster* CYP6G1, *N. lugens* CYP6AR1 and *Ae. aegypti* CYP6BB2. These point-mutations were present at low frequency in the Tiassalé-S line (∼1.6%) but strongly enriched in the Clothia-R line (∼97%) while their frequency remained low in other selected lines. Comparing the best *in silico* models obtained for the Tiassalé-S and Clothia-R CYP6M1 variants docked with clothianidin allowed identifying residues most likely interacting with the ligand in the active site (Figure 6D). This analysis suggested a 17% increase in the volume of the binding pocket in the Clothia-R variant (869 Å^3^) as compared to the Tiassalé-S variant (738 Å^3^). The presence of a hydrogen bound between clothianidin and the Val-369 (distance of 3.6 Å) was predicted in both variants though another hydrogen bound with Leu-485 was predicted in the Clothia-R variant due the rotation of clothianidin in the active site. This rotation also led to a lower distance between the Asp-215 and clothianidin suggesting a potential interaction of this residue with the ligand in the Clothia-R variant. This rotation also led to a different positioning of the nitro- and methyl-groups of clothianidin though the nitroguanidine moiety was still facing the heme, suggesting that preferred metabolites could be desmethyl-clothianidin, desmethyl-denitro-clothianidin or clothianidin-urea [28,29]. Overall, although the amino acid changes identified in the SRS regions between Tiassalé-S and Clothia-R variants did not significantly differed in their physio-chemical properties, *in silico* models suggest that they may affect the positioning of clothianidin within CYP6M1 active site and subsequent enzyme-ligand interactions.

**Figure 6.**
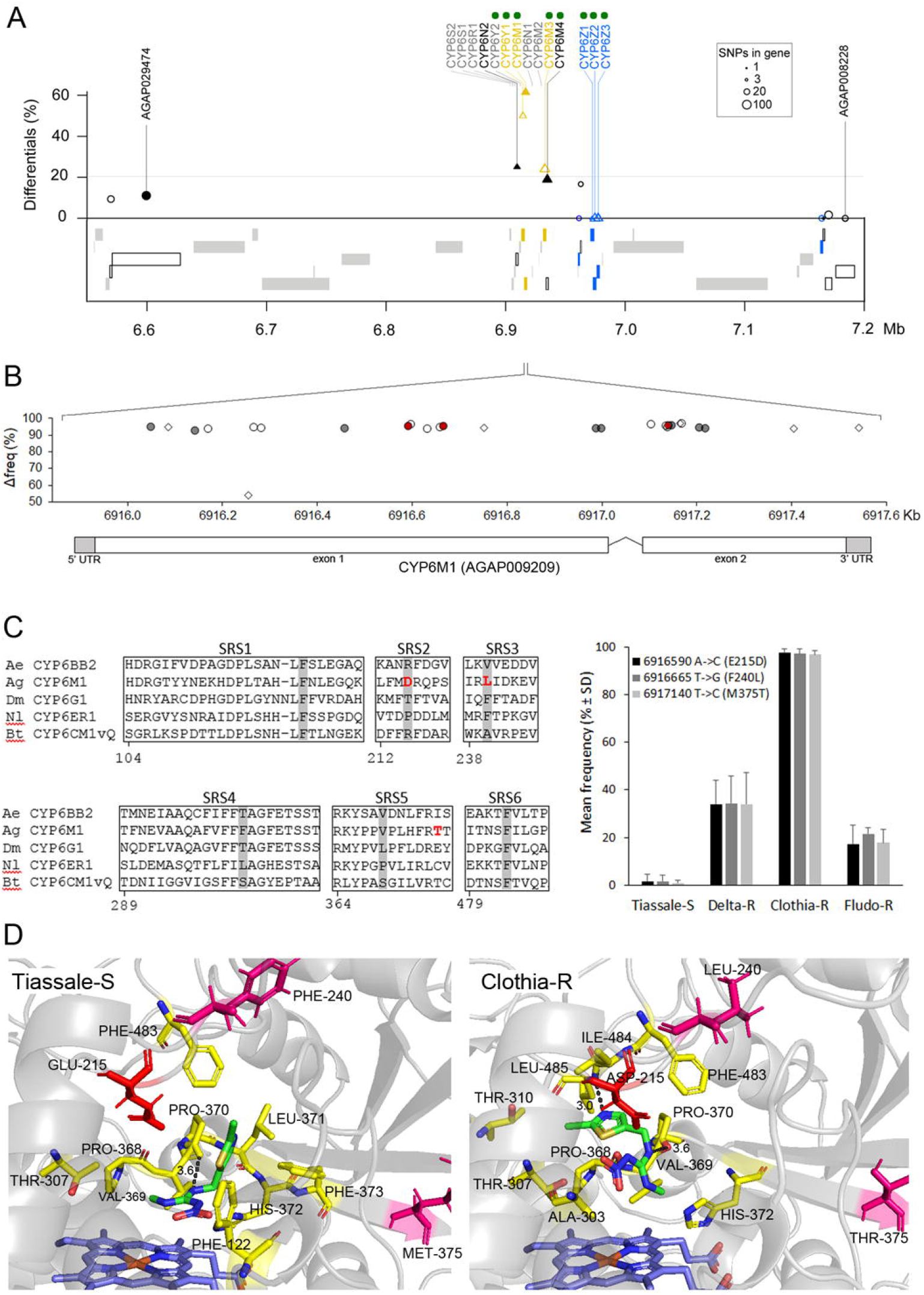
Response to clothianidin selection on the CYP6Ms locus. **A:** Overview of RNA-seq data at the CYP6M locus for the Clothia-R line. The Y axis shows the proportion of differential SNPs per gene as obtained by the frequency-based approach. Genes in grey were not covered by polymorphic SNPs. Significant transcription level variations in the Clothia-R line as compared to the non-selected Tiassalé-S line are indicated by colours (yellow: over-transcribed, blue: under-transcribed, black: not differentially transcribed). Triangles indicate candidate genes. Filled symbols indicate the presence of differential/outlier SNPs affecting protein sequence. Green dots indicate genes showing high protein homology with genes conferring resistance to neonicotinoids in other insect species. **B:** Distribution of differential SNPs on *CYP6M1*. The Y axis shows the frequency variation between the Clothia-R line and the Tiassalé-S line. Empty symbols: synonymous variations, filled symbols: non-synonymous variations. Symbol form indicate that the variant (circles) or the reference (lozenges) allele is enriched in the Clothia-R line. Red symbols indicate the two non-synonymous variations affecting the conformation of the active site. **C:** Focus on non-synonymous variations affecting the active site. Left: non-synonymous variations affecting CYP6M1 substrate recognition site regions (SRS) in the Clothia-R variant. Amino acid in grey are those shown to interact with the binding of the neonicotinoid imidacloprid in *Bemiscia tabaci* CYP6CM1vQ (Karunker et al 2009), *Nilaparvata lugens* CYP6ER1 (Pang et al 2006) and *Aedes aegypti* CYP6BB2 (Riaz et al 2012). Right: allele frequencies of the two non-synonymous variations likely interacting with the binding of clothianidin across all lines. **D:** In silico models of CYP6M1 protein variants identified in the Tiassalé-S and Clothia-R lines docked with heme (blue) and clothianidin (green). The best models obtained according to Rosetta-computed force fields are shown for each variant. Yellow residues are within 3.5 Å of the ligand. Residues differing between the two variants in SRS regions are shown in magenta. The Glu/Asp-215 located within 3.5 Å of the ligand in the Clothia-R variant but not in the Tiassalé-S variant is shown in red. Dark dashed lines show hydrogen bonds between the ligand: Val-369 (both variants), Leu-485 (Clothia-R variant only).

The association between *CYP6M1, CYP6M2* and *CYP6M3* transcription levels and clothianidin resistance was then examined at generation G31 using RT-qPCR. The transcription level of each gene was compared between the Tiassalé-S and the Clothia-R lines in combination with pre-exposure to the P450 inhibitor piperonyl butoxide (PBO) and subsequent exposure to clothianidin (Additional file 6). Among the 3 *CYP6M*s, only *CYP6M1* showed an over-transcription in the Clothia-R line as compared to the Tiassalé-S line while *CYP6M2* and *CYP6M3* were under-transcribed. Though the over-transcription of *CYP6M1* was not significant (P=0.07), it became significant in clothianidin survivors (P=0.0014) suggesting an association between its over-expression and clothianidin survival. Exposing Clothia-R individuals to 4% PBO alone did not significantly induce *CYP6M1* transcription (P=0.19). Pre-exposing Clothia-R individuals to PBO before significantly decreased clothianidin survival by 20% (P<0.05). such exposure also restaured *CYP6M1* transcription level in survivors to a comparable level as in unexposed individuals suggesting a lower survival of individuals over-transcribing CYP6M1 following its inhibition by PBO.

## Discussion

The pyrethroid-neonicotinoid combination Fludora® Fusion was pre-qualified for IRS usage by the World Health Organization in December 2018 [30]. Field trials performed in multiple African countries confirmed the long lasting effect of this combination on various sprayed surfaces and its good efficacy against pyrethroid-resistant malaria vectors [25,31]. Though resistance risk assessment is not mandatory in the public health insecticides evaluation scheme, understanding the resistance dynamics of novel insecticide-based product is essential for anticipating resistance management strategies and extend their lifespan. In this context, the aim of the present study was to investigate through a controlled-selection experiment the potential of the African malaria vector *An. gambiae* to develop resistance this novel insecticide combination across multiple generations and investigate associated mechanisms. Because the operational advantage of this novel product essentially stands on the association of two active ingredients with distinct modes of action, our controlled selection experiment compared resistance dynamics to each insecticide separately and to their combination. As neonicotinoid resistance alleles may already be circulating in mosquito populations indirectly exposed to neonicotinoids used in agriculture [26,27], a field-derived *An. gambiae* line originating from the agricultural area of Tiassalé, Côte d’Ivoire was used as a parental line selection.

### Fludora® Fusion mixture and its components select different adaptive responses

Previous studies showed that *An. gambiae* from the Tiassalé area (Côte d’Ivoire) are highly resistant to the pyrethroid deltamethrin, the carbamate bendiocarb, the organochlorine DDT and moderately resistant to the organophosphate fenitrothion [32]. This multi-resistance phenotype was associated with a high frequency of the Kdr west L1014F mutation (∼80%), a moderate frequency of the ace1 G119S mutation (∼50%) and the over-transcription of multiple P450s previously associated with insecticide resistance [33]. Bioassays performed on the Tiassalé-S line confirmed its lower resistance to insecticides following the introgression of susceptible alleles. This was confirmed by its lower frequencies for the *Kdr west* L1014F and the Ace1 G119S mutations with most individuals carrying them being heterozygotes. As predicted by the presence of pyrethroid-resistance alleles in the Tiassalé-S line, deltamethrin selection rapidly led to an increased resistance to this insecticide in the Delta-R line. This resistance phenotype was tightly associated with the increased frequency of the *Kdr west* mutation, confirming the key role of this mutation in deltamethrin resistance [9,12]. Only a few detoxification enzymes were over-transcribed in the Delta-R line excluding key pyrethroid metabolizers such as *CYP6M2* and *CYP6P3* previously identified in Tiassalé [15,33] or any other detoxification enzyme commonly associated with insecticide resistance [14,34]. Such absence of key metabolic enzymes in the Delta-R line suggests that the selection pressure applied (>50% mortality) rather selected for individuals carrying the *Kdr west* mutation than those carrying metabolic resistance alleles. Hence, the rapid selection of this major mutation could have prevented the selection of metabolic resistance alleles of lower importance in deltamethrin resistance [35]. This was supported by polymorphism data showing no selection signature at metabolic resistance *loci* in the Delta-R line. Overall, this supports that the selection of target site resistance mechanisms are favored under intense selection pressure [36,37]. No such increased resistance was observed with Fludora® Fusion mixture after 33 generations, suggesting that in our experimental conditions, the clothianidin-deltamethrin combination prevented the selection of resistance alleles having a significant impact on the overall resistance phenotype. The gradual decreased frequency of the *Kdr west* mutation in the Fludo-R line also supports the role of clothianidin in precluding its selection by the deltamethrin-clothianidin mixture.

However, we showed that selection with the neonicotinoid clothianidin alone led to an increased resistance to this insecticide although the resistance level was moderate and remained stable. RNA-seq data generated from the Clothia-R line suggests that detoxification genes previously associated with insecticide resistance were affected by clothianidin selection. This included multiple P450s from *CYP6* and *CYP12* families frequently associated with insecticide resistance. Though drift may have occurred during the selection process, the impact of clothianidin selection on detoxification enzymes associated with resistance was supported by the strong selection signature observed for the Clothia-R line at the *CYP6M* locus. Indeed, multiple *CYP6* genes located at this locus showed high protein orthology with P450s conferring resistance to neonicotinoids in other insect species, including *Ae. aegypti CYP6BB2* [38], *D. melanogaster CYP6G1, CYP12D1* and *CYP6A8* [39–41] and *B. tabaci CYP6CM1vQ* [42]. Focusing on this locus revealed that *CYP6M1* showed the strongest selection signature and was affected by multiple non-synonymous mutations with two of them (E215D and F240L) located at positions previously suggested to interact with the binding of the neonicotinoid imidacloprid in *B. tabaci* CYP6CM1vQ, *D. melanogaster* CYP6G1, *N. lugens* CYP6AR1 and *Ae. aegypti* CYP6BB2 [38–42]. *In silico* modelling of CYP6M1 variants suggested that structural differences associated with these amino acid changes may enhance clothianidin metabolism in the Clothia-R line. RT-qPCR validation confirmed the over-transcription of *CYP6M1* in the Clothia-R line although transcription ratios were lower than those inferred from RNA-seq. Nevertheless, *CYP6M1* transcription level was further augmented in Clothia-R individuals surviving clothianidin supporting its association with clothianidin resistance. This effect was lowered by PBO pre-exposure, suggesting that the biochemical inhibition of P450s led to a lower survival of individuals over expressing *CYP6M1*, which further supports its contribution to the resistance phenotype. Although this needs to be functionally validated, the present study suggests that both transcriptional regulation and variant selection of *CYP6M1* are associated with clothianidin resistance in *An. gambiae*. From a larger perspective, the present study supports the key role of P450s in neonicotinoids resistance as previously observed in other insect species [41,43–46].

In addition to detoxification enzymes, RNA-seq data pointed out the striking over-transcription of multiple nicotinic receptor subunits in the Delta-R and Clothia-R lines though no selection signatures were observed at these *loci*. No differential non-synonymous SNP was detected within these genes, invalidating the presence of neonicotinoid target-site mutations in these lines. It has been shown that the modification of synaptic Na+ homeostasis by 1014F Kdr mutation lead to a decrease of intracellular Ca^2+^ concentration which in turn affects synaptic transmission [47]. As neonicotinoids also affect intracellular Ca^2+^ concentration [48], it is probable that the over-expression of nicotinic receptors observed in the Delta-R and Clothia-R lines rather reflect physiological compensation mechanisms to altered synaptic transmission than direct resistance mechanisms. Finally, a significant increase of DDT resistance was observed in all selected lines as compared to the Tiassalé-S line. Though increased DDT resistance is likely associated with the *Kdr west* mutation in the Delta-R line, this is less likely for the Clothia-R and Fludo-R lines showing lower frequencies. However, genes encoding detoxification enzymes commonly associated with DDT resistance such as *GSTE2* [49], *CYP6Z1* [50] and *CYP6M2* [14,17,33] were not over-transcribed in these two lines, suggesting that the over-transcription of other resistance alleles might explain the observed phenotype. For instance, the over-transcription of *CYP4G16* in these lines in conjunction with multiple cuticle proteins is of interest as this particular P450 was associated with cuticle-based resistance in *An. gambiae* [51,52].

### Pyrethroid-neonicotinoid combination as a new tool for vector control in Africa

The use of the same insecticide classes targeting only two biochemical targets (the AchE and the VGSC) for decades has led to the selection and spread of insecticide resistance throughout Africa [53]. Insecticide resistance is now considered as a significant burden for malaria control with *Anopheles* populations often being resistant to multiple insecticide families [7]. Pyrethroid resistance is of major concern as these insecticides are mainly used for impregnating bednets [54,55]. In this context, the development of novel vector control products that can be used for controlling pyrethroid resistance has been encouraged by WHO [56]. Among them, IRS formulations combining two insecticides with different modes of action such as Fludora® Fusion are of high interest. Though this IRS combination proved to be efficient for several months against African malaria vectors [23–25], its long-term efficacy and its impact on insecticide resistance dynamics remains to be confirmed.

Although our controlled selection experiment was performed on a single field-derived *An gambiae* line and our results may have been impacted by drift effects, the absence of resistance development to Fludora® Fusion mixture suggest that clothianidin/deltamethrin combination represents an added value to current insecticides used for controlling malaria vectors in Africa. In particular, the potential of clothianidin to prevent the selection of *Kdr* mutations appears of high interest for managing pyrethroid resistance [57,58]. In turn, the selection of P450s associated with clothianidin resistance was impaired when deltamethrin was added, suggesting the added value of combining these two insecticides in delaying the emergence of neonicotinoid resistance. Such results are also in favor of the concomitant use of pyrethroid-impregnated bednets and neonicotinoid-based IRS products for vector control within the same location.

However, the present study also suggests that the use of clothianidin alone can rapidly select for metabolic resistance alleles in *An. gambiae*. P450-mediated cross-resistance between pyrethroids and neonicotinoids has previously been observed in various insect species [59–61] and our results support that this may also occur relatively rapidly in *An. gambiae*. Subsequent field studies may allow deciphering if these resistance alleles are only present in agricultural areas where neonicotinoids are regularly used or have already spread throughout Africa, and if they will be selected along the deployment of neonicotinoids for malaria control. In this frame, the candidate genes identified here provides the first set of neonicotinoid resistance markers in *An. gambiae*.

## Conclusions

The present study supports the potential of neonicotinoid-pyrethroid combination as a novel vector control tool having the ability to delay the selection of resistance. As such combination appears to show a good efficacy against pyrethroid resistant populations, their deployment within an integrated vector control framework and under careful monitoring may be of added value for managing insecticide resistance in African malaria vectors. Conversely, the deployment of neonicotinoids alone rises concerns as this may rapidly lead to the emergence and spread of neonicotinoid resistance alleles in malaria vectors as observed in other insect species.

## Methods

### Mosquitoes

*A. gambiae* larvae were collected in 2010 in the vicinity of Tiassalé in the south of Côte d’Ivoire, an intensive agricultural area which has high coverage of pyrethroid-impregnated mosquito nets together with an intensive use of pesticides for crop protection including pyrethroids, organophosphates, carbamates and neonicotinoids [62]. Previous studies showed that malaria vectors from this area carry multiple resistance alleles conferring resistance to various insecticide families including pyrethroids, DDT, carbamates and organophosphates [32,33,63]. After species identification, *An. gambiae sensus stricto* specimens were used to create the Tiassalé strain that was then kept without insecticide selection pressure at the Centre Suisse de Recherche Scientifique en Côte d’Ivoire (CSRS). In order to maximize the dynamic range of resistant allele frequency variations during the selection process, a moderately resistant line (Tiassalé-S line) created by mass-crossing the Tiassalé line once with the fully susceptible line Kisumu was used for the controlled selection experiment. The Tiassalé-S line was then maintained without selection pressure for two generations before selection. All the lines described thereafter were maintained in the LECA tropical insectaries under standard conditions (29°C, 90% relative humidity, 14h/10h light/dark period). Larvae were bred in deionized water and fed with TetraMin fish flakes. Adults were fed on filter papers impregnated with a 5% honey solution and blood feeding of adult females was performed on mice.

### Controlled selection

The Tiassalé-S line was divided into four distinct lines. The first line (Tiassalé-S) was kept without selection and used as control for the selection experiment. The three other lines were selected in parallel with the pyrethroid deltamethrin (Delta-R line), the neonicotinoid clothianidin (Clothia-R line) or a mixture of deltamethrin and clothianidin in equivalent proportion (8:1 w/w) to *Fludora® Fusion* (Fludo-R line). Because filter paper impregnation is not appropriate for clothianidin (crystallization of the active ingredient) all insecticide selections were performed using 250 mL glass bottles impregnated with insecticide. All insecticides used for bottle impregnation were obtained as pure active ingredients from Sigma. Deltamethrin was diluted in 100% acetone while clothianidin and clothianindin/deltamethrin mixture were diluted in an acetonic solution containing 17% (v/v) of mero solvent (MERO EC733, an emulsifiable concentrate containing 81.7% of rapeseed fatty acid esters in ethoxy-7-tridecanol). Bottles were impregnated with 1 mL of insecticide solution. For each line, insecticide selection consisted in introducing at least 15 batches of non-blood fed females into insecticide-impregnated bottles. Doses used for selection were initially calibrated in order to reach 60% mortality in the in the Tiassalé-S line as follows: deltamethrin 10 µg/mL; clothianidin 0.45 µg/mL; clothianindin/deltamethrin mixture 0.45/0.05625 µg/mL (w/w ratio of 8:1, thereafter designated as Fludora® Fusion mixture). Exposure times were as follows: deltamethrin 15 min, clothianidin 20 min, Fludora® Fusion mixture 15 min. These times were slightly adjusted through the selection process in order to maintain an equivalent selection pressure between lines (between 50-70% mortality). Mortality rates were recorded after a recovery time of 72h in order to consider the slower effect of clothianidin and Fludora® Fusion mixture. Survivors were then transferred into new cages and blood-fed to generate eggs for the next generation. Selection was performed for each line at each generation until G33 except for generations G3, G4, G8, G13, G15, G17, G20, G22, G27, G29 and G32 in order to limit drift effects as population size was low for one line.

### Resistance monitoring through the selection process

The resistance profile of the parental line Tiassalé-S to the four main insecticide classes was characterized before the start of the selection experiment (G0) using Standard WHO susceptibility assays. Insecticide exposures were performed according to WHO standard procedures [64] using test tubes equipped with filter papers impregnated with the following insecticides: deltamethrin 0.05%; DDT 4%; bendiocarb 0.5% and fenitrothion 1%. Exposure time was fixed to 1h for each insecticide. At least five lots of 20 five-day-old adult females were tested for each insecticide. Mortality rate was recorded after a 24h recovery time during which mosquitoes were provided a 5% honey solution. Mortality rates were expressed as mean mortality ± Wald’s confidence intervals.

The resistance level of each selected line to its respective insecticide was monitored at generations G0 (parental Tiassalé-S line), G2, G5, G7, G9, G11, G13, G17 and G33. Insecticide testing was performed using CDC bottles impregnated with a constant dose of insecticide corresponding to the initial dose used for selection (see above). Exposures were performed on 5 batches of 20 non-blood fed females per insecticide. Mortality rates were recorded after a 72h recovery time in order to consider the slower effect of clothianidin. The resistance level of the non-selected Tiassalé-S line to each insecticide was also monitored in order to account for variations across generations inherent to insecticide solution preparation. Mortality rates were expressed as mean mortality ± Wald’s confidence intervals. Differences between each selected line and the control line Tiassalé-S at each generation were tested using Fisher exact tests on mortality proportions (N=5). At G17, the resistance level of each line to the three insecticides were compared using the same methodology in order to assess cross resistance patterns between lines. The resistance levels of each line to DDT, bendiocarb and fenitrothion were also measured at G17 using standard WHO susceptibility assays as described above.

### Target sites mutations

The frequencies of target-site mutations previously identified in the Tiassalé strain (*Kdr west* L1014F and *Ace1* G119S) were monitored for each selected line through individual genotyping. Kdr L1014F mutation was genotyped at generations G0, G2, G7, G9, G11 and G17. Because the acetylcholinesterase was not targeted by any insecticide used for selection, the frequency of the ace1 G119S was only genotyped at G0 and G17. At each generation, genomic DNA was extracted from 30 adult females of each line using the cetyl-trimetylammonium bromide (CTAB) method [65]. Genomic DNA was resuspended in 20 µL nuclease-free water, quantified using the Qubit DNA BR assay (Thermofisher Scientific) and diluted to 0.5 ng/µL for genotyping. The Kdr L1014F and Ace1 G119S mutations were genotyped using the TaqMan qPCR methods described in [66]. Quantitative PCR reactions were performed on a CFX96 Real Time system (Bio-Rad) with PCR cycles as follows: 95°C for 10 min, followed by 40 cycles of 95°C for 10 sec and 60°C for 45 sec. For each mutation, individuals were scored as homozygous susceptible/resistant or heterozygous based on the intensity of the HEX/FAM channels at the end of the PCR reaction as compared to positive and negative samples of known genotypes.

### RNA Library preparation and sequencing

Differential gene expression between each selected line and the non-selected line Tiassalé-S was investigated at the whole transcriptome level at G17 using RNA-sequencing (RNA-seq). For each line, four pools of 30 three-day-old non-blood fed females not previously exposed to insecticide were used. Total RNA was extracted from each pool separately using Trizol (Life Technologies) according to manufacturer’s instructions. Total RNA was then treated with DNase to remove genomic DNA contaminants. RNA-seq libraries were prepared from 150 ng total RNA using NEBNext® Ultra™ II Directional RNA library Prep Kit for Illumina (New England Biolabs) following manufacturer’s instructions. Libraries were quantified using the Qubit DNA BR assay (Thermofisher Scientific) and quality checked on a Bioanalyzer (Agilent). Libraries were sequenced in multiplex as single 75 bp reads on a NextSeq 500 sequencer (Illumina) by Helixio (Clermont-Ferrand, France). After unplexing and quality check using FastQC, reads were loaded into Strand NGS V 3.2 (Strand Life Sciences) and mapped against the AgamP4 assembly and AgamP4.12 geneset using standard parameters (min identity 90%, max gaps 5%, min aligned length 35 bp, ignore reads with more than 5 matches, trim 3’ ends of reads with average quality ≤ 20). Mapped reads were then filtered based on their sequence quality and mapping quality as follows: Mean read quality ≥ 20, number of N ≤ 5, alignment score ≥ 90, mapping quality ≥ 120, number of matches = 1. The remaining reads (∼90% of sequenced reads) were used for subsequent analyses.

### Differential gene transcription analysis

Differential Transcription analysis was performed on all protein coding genes with normalization and quantification based on the DE-Seq algorithm [67]. Only the 10829 genes showing a coverage ≥ 4 reads/kb in all replicates of all conditions were kept. Transcription levels between each selected line and the parental line Tiassalé-S were then compared across the four biological replicates using an ANOVA followed by a Tukey HSD test. P values were adjusted for multiple testing corrections using the Benjamini-Hochberg method [68]. Genes showing a transcription ratio ≥ 1.5 fold in either direction and a P value ≤ 0.005 in any selected line as compared to the Tiassalé-S line were considered differentially transcribed following insecticide selection.

### Gene ontology terms enrichment

For each line, genes significantly over- and under-transcribed were subjected to a Gene Ontology term (GO-term) enrichment analysis using the functional annotation tool DAVID (http://david.abcc.ncifcrf.gov, [69]). Reference gene list consisted in the 10829 genes detected by RNA-seq. For each line, over- and under-expressed genes were considered separately and GO-terms showing a Fisher’s Exact test P value< 0.05 were considered enriched as compared to the reference list.

### Focus on resistance genes

Heat maps reflecting transcription profiles of candidate genes (*i*.*e*. detoxification enzymes including cytochrome P450s, carboxy/cholinesterases; transferases; ABC-transporters; cuticle proteins; redox enzymes and nervous receptors) across all lines were generated using TM4 MeV [70]. Genes previously associated with resistance to insecticides used in vector control were identified based on existing literature [15,53,71–72]. Because neonicotinoid resistance mechanisms have been poorly investigated in *An. gambiae*, orthologous genes to those conferring neonicotinoid resistance in other insect species were identified by protein sequence homology using NCBI BlastP against AgamP4 proteins with default parameters. Only *An. gambiae* genes showing a protein homology score ≥ 300 (E value ≤ 1E-95) to known neonicotinoid resistance genes were retained (Additional file 7).

### Polymorphism calling

Polymorphisms were called using strand NGS V 3.2 against all protein-coding genes of the AgamP4 assembly using standard parameters (ignore homopolymer stretches greater than 4 bp and adjacent positions, coverage ≥ 30 and ≤ 5000, reads supporting the variant allele ≥ 2, base quality ≥ 20, variant confidence score ≥ 200 and strand bias ≤ 25). Among variations passing these filters, only substitutions and indels were retained for further analyses. This calling and filtering strategy allowed detecting 166220 polymorphisms across all lines. The following genic effects were then computed based on AgamP4.12 annotation: synonymous coding, non-synonymous coding, start-lost, stop gained, stop-lost, frameshift coding, splice site, essential splice site, 5’ UTR, 3’ UTR, upstream (within 1500 bp of gene start), downstream (within 1500 bp of gene stop), near gene (within 100 bp of gene).

### Selection signatures associated with insecticide selection

Selection signatures associated with insecticide selection were investigated using the 145008 bi-allelic SNPs (substitutions or indels) that were polymorphic (i.e. showing ≥ 5% allele frequency variation between the Tiassalé-S line and any selected line). A first approach consisted in comparing the mean allele frequency of each SNP between each selected line and the Tiassalé-S line across the four replicates using a Student’s test followed with a Benjamini and Hochberg multiple testing correction [68]. SNPs showing a mean frequency variation between any selected line and the Tiassalé-S line ≥ 50% in either direction and a corrected P value ≤ 0.001 were considered as associated with insecticide selection (Differential SNPs). A second approach consisted in assessing F_ST_ departure from neutrality using the Bayesian method implemented in BayeScan version 2.1 [73]. A separated analysis was performed for each selected line consisting in contrasting the selected line versus the Tiassalé-S line across all replicates. Default settings were used except that prior odd was set to 1000 in order to increase stringency. Genes showing a Bayscan Q□value of zero were considered as ‘Outliers’. For each selected line, the proportions of ‘Differential SNPs’ and ‘Outlier SNPs’ per gene were computed and plotted along chromosomes using gene centers as genomic coordinates.

### Impact of clothianidin selection on the CYP6M locus

The association of *CYP6M1, CYP6M2* and *CYP6M3* with clothianidin resistance were further studied at generation G31 using RT-qPCR. These genes were chosen based on their RNA-seq expression profile, selection signature and their potential role in insecticide resistance. For each gene the following conditions were compared: Tiassalé-S unexposed (Tiassalé-S); Clothia-R unexposed (Clothia-R); Clothia-R surviving clothianidin exposure (Clothia-R surv); Clothia-R exposed to the P450 inhibitor piperonyl butoxide PBO (Clothia-R PBO+); Clothia-R pre-exposed to PBO and surviving clothianidin exposure (Clothia-R PBO+ surv). PBO exposure consisted in exposing batches of 25 mosquitoes to 4% PBO for 1h in glass bottles as described in [74]. Clothianidin exposure consisted in exposing batches of 25 mosquitoes for 1h to the same dose of clothianidin used for selection (0.45 µg/mL). Mosquitoes from all conditions were sampled at 72h post clothianidin exposure (6-days old). Such recovery time allowed considering the slow effect of clothianidin together with minimizing gene induction/repression effects that may follow insecticide exposure. For each condition, total RNA was extracted from four pools of 12 to 20 mosquitoes using Trizol (Life Technologies) according to manufacturer’s instructions. Two µg total RNA were treated with DNase I (Invitrogen) and reverse transcribed using superscript III (Invitrogen) and Oligo DT_20_ primer according to manufacturer’s instructions. cDNA samples were then diluted to 1/100 for qPCR amplification. Each sample was amplified as three PCR replicates. Quantitative PCR reactions were performed on a CFX qPCR system (Bio-Rad). Each qPCR reaction contained 12.5 µL iQ SYBR Green Supermix (Bio-Rad), 0.75 µL of each primer (10 µM each), 6 µL nuclease-free water and 5 µL diluted cDNA template. PCR cycles were as follows: 95°C 3 min followed by 40 cycles consisting of 95°C for 15 secs and 30 secs at 60°C. Specific primer pairs were used for each gene (Additional file 8) and amplification specificity was verified by melt curve analysis. Data analysis was performed according to the ΔΔ_Ct_ method taking into account PCR efficiency [75] using the gene encoding the ribosomal protein S7 (AGAP010592) as control. Results were expressed as mean transcription fold change (± SD) across the four biological replicates as compared to unexposed Tiassalé-S individuals. Differences across conditions were tested using an ANOVA across all conditions followed by Fischer tests between pairs of conditions (N=4).

Differential SNPs impacting *CYP6M1* (AGAP008209) were further examined. Amino acid changes resulting from differential non-synonymous SNPs were mapped to CYP6M1 protein sequence and aligned using ClutalW against the protein sequence of four *CYP6*s known to metabolize neonicotinoids: *Drosophila melanogaster* CYP6G1[39–41], *Bemiscia tabaci* CYP6CM1vQ [42], *Nilaparvata lugens* CYP6ER1[76] and *Aedes aegypti* CYP6BB2 [38]. P450 substrate recognition sites (SRS) were identified from protein alignment as defined from CYP3A4 structure [77]. The impact of amino acid changes selected by clothianidin on CYP6M1 structure and clothiainidin docking was then investigated by comparing the protein models obtained for the two distinct CYP6M1 variants identified in the Tiassalé-S and Clothia-R lines. For each variant, the amino acid sequence used for protein modelling differed by the 11 amino acid changes showing a differential frequency higher than 50% (Additional file 9). CYP6M1 protein models were computed by the SWISS-MODEL server using the crystallized human P450 CYP3A4 as template (protein Accession 5VCD). The Rosetta algorithm was then used to refine protein structure and to compute the most probable positions for the heme and clothianidin in the active site of each variant according to Rosetta-computed force fields [78]. The volume of the binding pockets of each variant was estimated using POVME 2.0 [79].

## Declarations

### Ethics approval and consent to participate

Blood feeding of adult mosquitoes was performed on mice. Mice were maintained in the animal house of the federative structure Environmental and Systems Biology (BEeSy) of Grenoble-Alpes University agreed by the French Ministry of animal welfare (agreement n° B 38 421 10 001) and used in accordance to European Union laws (directive 2010/63/UE). The use of animals for this study was approved by the ethic committee ComEth Grenoble-C2EA-12 mandated by the French Ministry of higher Education and Research (MENESR).

## Consent for publication

Not applicable.

## Availability of data and materials

RNA-seq sequence data reported in this study have been deposited to the European Nucleotide Archive (ENA; http://www.ebi.ac.uk/ena) under the accession numbers PRJEB44777. The other datasets used and/or analyzed during the current study are available as supplementary information and/or from the corresponding author on reasonable request.

## Authors’ contributions

MGZ contributed to study design, conducted experiments, contributed to data analysis and drafted the manuscript. JMB contributed to RNA-seq data analysis and contributed to manuscript writing. JT provided technical help for mosquito rearing, bioassays and molecular work. FL provided technical help for molecular work. MC, CS and BKF provided the initial Tiassalé line and contributed to experiment design. JM, FS and SH contributed to study design and funding acquisition. SR and JPD conceived the study, analyzed data and wrote the manuscript. JPD coordinated the study.

## Acknowledgements

We thank the Centre Suisse de la Recherche Scientifique (CSRS) de Côte d’Ivoire for providing the Tiassalé mosquito line. We thank Fabrice Chandre, Pierrick Labbé, Muriel Raveton, Frederic Boyer and Vincent Corbel for useful comments on the study design and the interpretation of results. We also thank Thierry Gaude for providing technical assistance in mosquito rearing and blood feeding.

## Supplementary information

**Additional file 1**. This .tif file describes the resistance of the parental line Tiassalé-S to insecticides commonly used for vector control. Insecticide susceptibility tests were performed using WHO test tubes equipped with papers impregnated with 0.05% deltamethrin, 0.5% bendiocarb, 1% fenitrothion and 4% DDT. Mortality rates are expressed as mean mortality ± 95% Wald confidence interval.

**Additional file 2**. This .xlsx table describes transcription data obtained for all genes detected by RNA-seq.

**Additional file 3**. This .tif file provides an overview of genes differentially transcribed in the three selected lines. The number of genes is indicated for each line. Numbers within bracket refer to candidate genes potentially involved in insecticide resistance.

**Additional file 4**. This .tiff document provides an overview of GO term enrichment analysis. Functional pathways enrichment analyses were based on genes significantly over- and under-transcribed in each selected line as compared to Tiassalé-S line using DAVID functional annotation tool (modified Fisher’s exact test with P < 0.05). Only GO terms from the ≪ biological process ≫ family showing an enrichment associated with a P value <0.05 and a minimum number of 9 genes are shown. Fold-enrichment (x axis), P value (color scale) and class size (dote size) are indicated.

**Additional file 5**. This .xlsx table describes SNPs data obtained from RNA-seq. Only genes containing polymorphic SNPs are shown.

**Additional file 6**. This .tif document shows the results of the association of *CYP6M* genes transcription levels with clothianidin resistance. A: RT-qPCR profiles of *CYP6* genes in the following conditions: Tiassalé-S unexposed (Tiassalé-S); Clothia-R unexposed (Clothia-R); Clothia-R surviving to clothianidin exposure (Clothia-R Surv); Clothia-R exposed to PBO (Clothia-R PBO+); Clothia-R exposed to PBO and surviving to clothianidin exposure (Clothia-R PBO+ Surv). Mosquitoes from all conditions were sampled at the same time corresponding to 72h after clothianidin exposure (6-days old). RT-qPCR was performed on four biological replicates per condition. Letters indicate significance between pairs of conditions (P<0.05). B: Effect of PBO pre-exposure on the survival of Clothia-R individuals to clothianidin. Mosquitoes were exposed to 4% PBO for 1h prior exposure to 0.45 µg/mL clothianidin for 1h. Mortality was recorded 72h after exposure. Letters indicate significance between conditions (P<0.05).

**Additional file 7**. This .xlsx table describes *An. gambiae* genes showing high protein homology with gene conferring resistance to neonicotinoids in other insect species.

**Additional file 8**. This .xlsx table describes primers used for RT-qPCR.

**Additional file 9**. This .docx document describes the amino acid sequence of the two CYP6M1 variants identified in the Tiassalé-S and Clothia-R lines used for protein modelling.

