## Supplementary figures and images for "Neonicotinoid and pyrethroid combination: A tool to manage insecticide resistance in malaria vectors? Insights from experimental evolution"

### Additional file 1

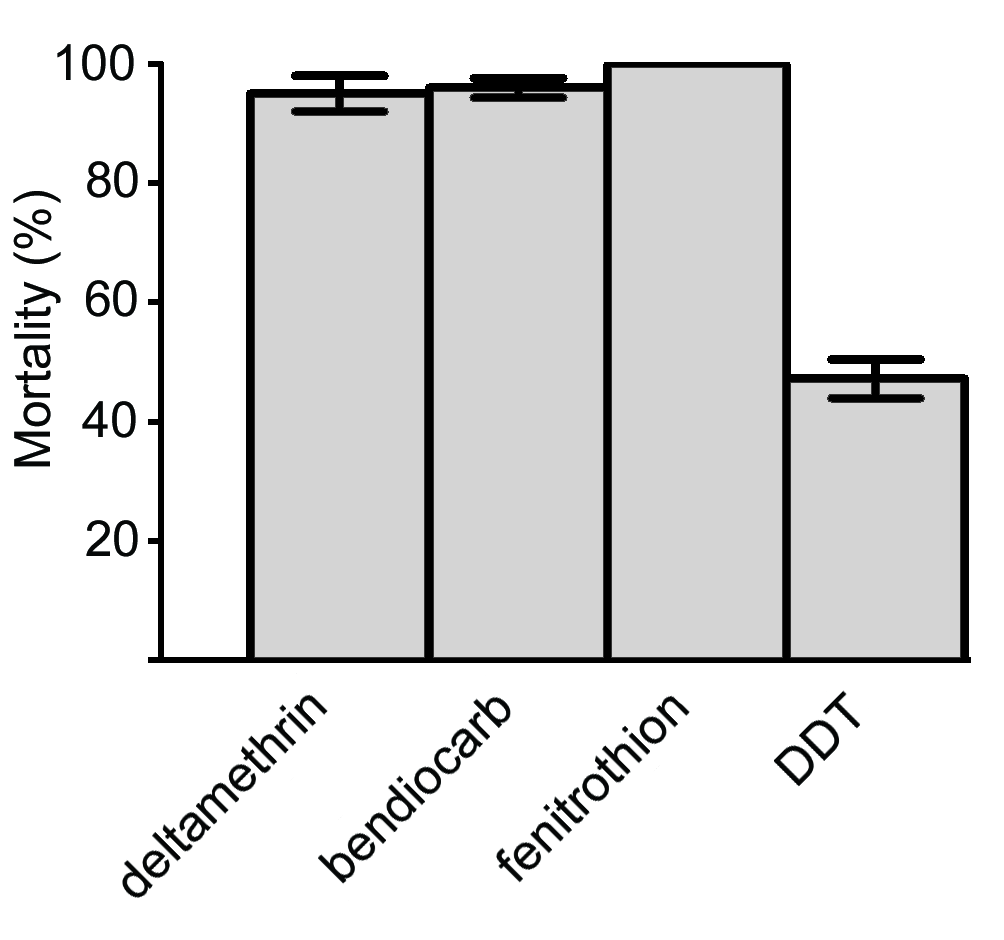

### Additional file 3

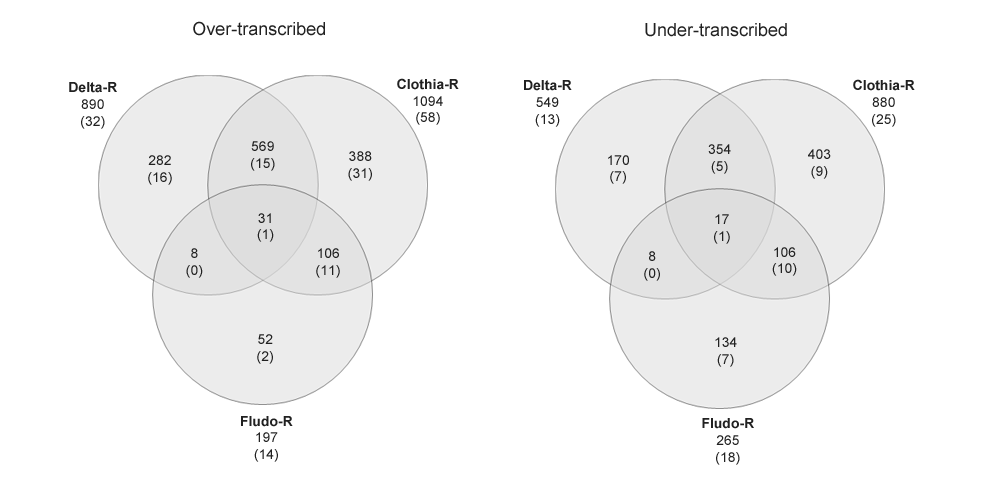

### Additional file 4

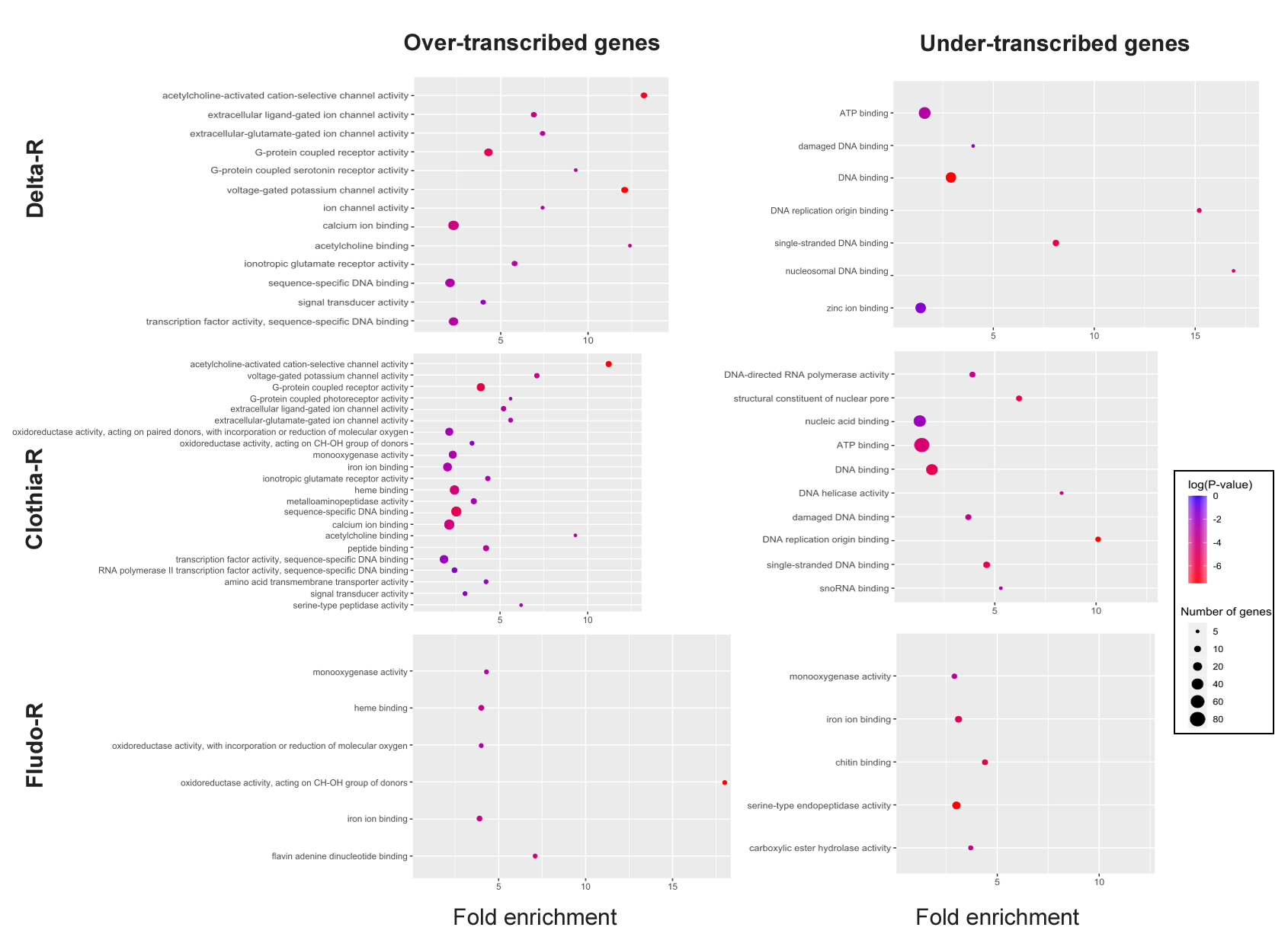

### Additional file 6

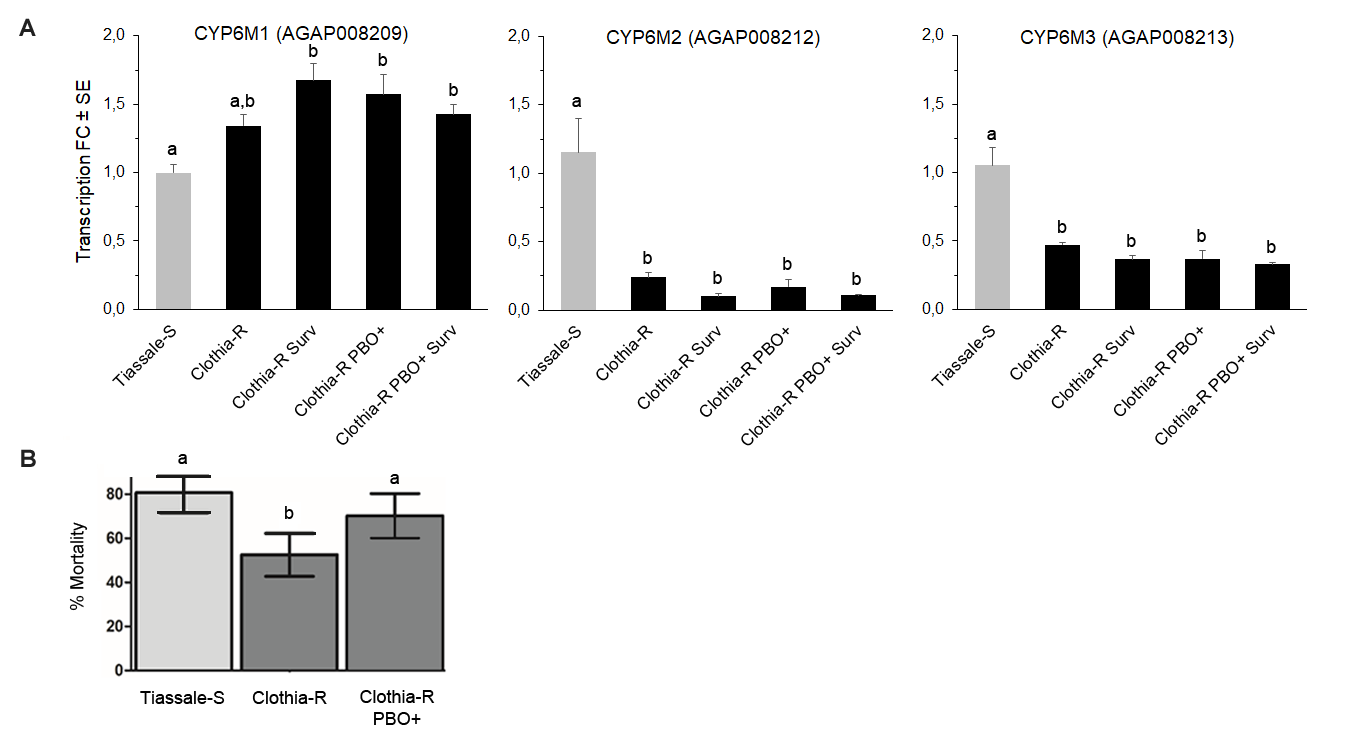
