## Additional file 9 for "Neonicotinoid and pyrethroid combination: A tool to manage insecticide resistance in malaria vectors? Insights from experimental evolution"

**Additional file 9.** Amino acid sequence of the two CYP6M1 variants identified in the Tiassale-S and Clothia-R lines used for protein modelling. For each variant, the amino acids in grey boxes differed by more than 50% frequency as compared to the reference genome AgamP4.12 sequence (AGAP008209). Amino acids in red are those supported by an allele frequency variation >50% between the Tiassalé-S line and the Clothia-R line.

**An. gambiae CYP6M1 (Tiassalé-S line)**

MWFPTIEVLVALLALLGGAVYFIVRKQSYWKERG**V**PHPKPTFFFGSFKDAGTKIHFTEEVERHY**AI**YKGKHPFIGVY**M**LTTPVVLPLDLELIKAIFVKDFQYFHDRGTYYNEKHDPLTAHLFNLEGQKWRNLRNKMTPTFTSGKMKMMFPTVVAAGQQLRDFMEENVQKH**G**EMELKDVMARYTTDVIGTCAFGIECNSMRDPDAEFRAMGKLFM**E**RQPSQFVNMMVQFSPKLSRLLGIR**F**IDKEVS**A**FFLKVVRDTIDYRV**K**NGIQRNDFMDLMIRMLQNTENPEEALTFNEVAAQAFVFFFAGFETSSTLLTWTLYELALNPEVQEKGRQCVQEVLAKHNGEMTY**D**AIH**D**MKYLDQILKESLRKYPPVPLHFR**M**T**A**QNYRVPDTDSVIEAGTML**F**IPIF**S**IQRDASLFPEPEKFDPERFSAEEEAKRHPFAWTPFGEGPRVCIGLRFGMMQARIGLAYLLQGFSFAPYEKTSIPMKFITNSFILGPREGLWLKVNKLESKQG

**An. gambiae CYP6M1 (Clothia-R line)**

MWFPTIEVLVALLALLGGAVYFIVRKQSYWKERG**I**PHPKPTFFFGSFKDAGTKIHFTEEVERHY**AL**YKGKHPFIGVY**L**LTTPVVLPLDLELIKAIFVKDFQYFHDRGTYYNEKHDPLTAHLFNLEGQKWRNLRNKMTPTFTSGKMKMMFPTVVAAGQQLRDFMEENVQKH**D**EMELKDVMARYTTDVIGTCAFGIECNSMRDPDAEFRAMGKLFM**D**RQPSQFVNMMVQFSPKLSRLLGIR**L**IDKEVS**A**FFLKVVRDTIDYRV**K**NGIQRNDFMDLMIRMLQNTENPEEALTFNEVAAQAFVFFFAGFETSSTLLTWTLYELALNPEVQEKGRQCVQEVLAKHNGEMTY**E**AIH**E**MKYLDQILKESLRKYPPVPLHFR**T**T**S**QNYRVPDTDSVIEAGTML**L**IPIF**A**IQRDASLFPEPEKFDPERFSAEEEAKRHPFAWTPFGEGPRVCIGLRFGMMQARIGLAYLLQGFSFAPYEKTSIPMKFITNSFILGPREGLWLKVNKLESKQG
